# Physiological tolerance of the early life history stages of fresh water prawn (*Macrobrachium rosenbergii* De Man, 1879) to environmental stress

**DOI:** 10.1101/159244

**Authors:** Jojy John, Vinu S Siva, Amit Kumar, Vinitha Ebenezer, Philimon Raika, Umer Khalifa, Subramoniam Thanumalya, Prakash Sanjeevi

## Abstract

Global climate change is transforming life on earth, causing widespread effects on all ecosystems. Among marine ecosystems, estuaries are considered as nursery grounds for marine and fresh water species. *M. rosenbergii*, a euryhaline species, migrate to the estuaries for breeding and spawning. The subsequent larval rearing takes place by experiencing variations in temperature and salinity conditions. The present study examines the effect of different temperature and salinity on the larval development and survival by observations on stored yolk utilization, cardiac performance, as well as changes in the rate of growth in body appendages and larval activity. The larvae showed 100% mortality at higher temperature (33.5 ± 0.5 °C) in all the salinity conditions (12 PPT, 15 PPT, and 20 PPT). The survival rate varied between 76-96 % on exposure to lesser temperature conditions. Likewise, the post-embryonic yolk lasted for 4 days at ambient temperature (29 °C); whereas, at 33.5 ± 0.5 °C, it lasted only for 2-3 days. There was an increase in total length of larvae, when exposed to higher temperature and salinity, independently or in combination, but at 33.5 ± 0.5 °C under all salinity conditions the larvae died on the 5^th^ day. For the cardiac performance, larval heart beat (*f*_H_) significantly increased for higher temperature and salinity conditions (20 PPT; 33.5 °C) and lowered at ambient condition 12 PPT; 29°C. Larval stroke volume *V*_s_, and Cardiac output Q□ were higher in ambient conditions and lowest in higher temperature and salinity conditions. However, temperature and salinity together did not show any significant effect on cardiac performance. On the other hand, the larval activity decreased significantly at higher temperature and salinity conditions, compared to ambient conditions but the interactive effect did not show any change. Thus, the physiological responses to temperature and salinity by the early life stages of *M. rosenbergii* could restrain the tolerance capability of the organism, thereby interfering in the successful completion of the larval development under the altered climatic conditions.

## INTRODUCTION

Earth’s climate is changing at a rapid pace, mainly because of the increased carbon dioxide emission caused due to anthropogenic activities (Solomon et al., 2009). Though climate change is a global phenomenon, its effects on living organisms manifest at very local levels (Helmuth, 2009), and the magnitudes of these changes/effects could considerably fluctuate from location to location. Estuaries are one such ecosystem that is influenced by a variety of anthropogenic stressors (Donders et al., 2008). Estuaries act as a natural shelter for all myriad forms of aquatic life on earth where the spawning and feeding of early life forms of fish and shellfish happen (Beck et al., 2001). The diversity, distribution and biological functions of the organisms living in estuaries are influenced by climate change stressors (Kinne, 1971; Pörtner, 2008; Pörtner, 2005; Portner and Knust, 2007; Widdicombe and Spicer, 2008). Climate change stressors including, rise in temperature, precipitation, salinity changes, sea level rise and ocean acidification pose deleterious impact on marine organisms and ecosystems (Brierley and Kingsford, 2009)

The giant fresh water prawn, *Macrobrachium rosenbergii*, is an indigenous species to south and south-east Asia (Holthuis, 1980). Lately, this prawn has been introduced to several other countries as commercially important aquaculture species (New, 2002). In their natural environment, *M. rosenbergii* is inhabited in various environments including fresh water streams, estuarine waters and canals connected to the sea (Jalihal et al., 1993; Shokita, 1979; Tiwari, 1955). The life history of *M. rosenbergii* is amphidromous in nature. The adults spend their life in the fresh water; after spawning, brooders migrate to estuarine waters for hatching. The larval development takes place in estuarine water, and after settling down, it returns to the fresh water (Nandlal and Pickering, 2005; New, 2002). However, fresh water prawn culture in India increased steadily since 1999 reaching a peak output of 42,780 t in 2005, but then declined to 6,568 t in 2009–2010 due to poor seed and brood stock quality (Nair and Salin, 2012).

In general, elevated temperature and salinity variations affect the metabolic rate, calorific intake and energy budget of decapods (Anger, 2003). Temperature and salinity are the important abiotic factors that control the growth and development of decapod crustaceans (Anger, 2003; Kinne, 1964; Kinne, 1971; Chand et al., 2015; Habashy and Hassan, 2010). Salinity around 12-15 PPT and temperature range from 28 to 30 °C appeared to be optimal for the adults and larvae (Ling, 1978). Salinity plays a critical role on egg, embryo and larval development during the life cycle of *M. rosenbergii.* Yen and Bart (2008) studied the negative effects of elevated salinity on the reproduction and growth of female *M. rosenbergii.* Salinity influences all aspects of larval biology including survival, development, morphology, the moulting cycle, growth, feeding, metabolism, energy partitioning, and behavior (Anger, 2003). Likewise, Guest and Durocher (1979) reported the necessity of brackish water for the completion of larval development in *M. amazonicum*.

Temperature, the other major factor, influences the species distribution, range of thermal tolerance and acclimatization of ectotherm organisms (Schmidt-Nielsen, 1997). In tropical environment, ectotherms have narrow range of thermal tolerance due to lack of seasonality in this region and most of them are living at the verge of their maximum thermal limit, making them vulnerable under global warming scenarios (Sunday et al., 2012). In crayfish, significant difference on the gonad development and spawning were observed at different temperatures (Carmona-Osalde et al., 2004). Similarly, the effect of high temperature showed irregular patterns of egg development in *M. americanum* (Sainz-Hernández et al., 2016). The growth pattern of the *M. rosenbergii* adults also changed, as temperature increased from low to normal/optimum, with the growth declining at the higher temperature (Habashy and Hassan, 2010). Furthermore, synergistic effects in combination with one or more environmental variables (e.g., temperature and salinity) also play a key role in the ecological and geographical distribution of a species. Nelson et al. (1977) reported interactive effects of salinity and temperature on the metabolic rate of juveniles of *M. rosenbergii*.

The persistence or the failure of a population is determined by the successful completion of all larval stages (Byrne, 2011). Even though the impact of varying salinities and temperature have been extensively studied in adult and juveniles of *M. rosenbergii*, we have poor understanding on the physiological consequences of individual as well as interactive impacts of salinity and temperature on the early life stages of this species. Under the predicted climate change scenario, understanding the physiological constrains and energy cost for the completion of larva stages are vital to know the adaptive capability of successive population. Hence, in the present study, we used yolk utilization, cardiac performance, larval activity, growth as proxy to know the physiological fitness of the organism under future climate change condition. These data may give insight on the impact of climate change stress on early life history stages of this important aquaculture species.

## MATERIALS AND METHODS

### Animal collection and maintenance

The adult male and female of *M. rosenbergii* were procured from a fisherman based at Cuddalore, Tamil Nadu. The shrimps were transported to the demonstration hatchery at the Centre for Climate Change Studies, Sathyabama University, Chennai in an oxygen filled polyethylene bags. After transfer, the shrimps were acclimatized for 2-4 hours to the laboratory condition and shifted to 4×500 l fiber tank filled with the fresh water and fitted with biological filters. The water temperature was maintained at 29 °C and salinity at 0 PPT. The animals were fed three times a day with grated potatoes and commercial prawn feeds. Ten to 30% of the water was exchanged once in 3 days to maintain the quality.

### Experimental set up

Twenty spawned brooders were kept in 4×300 liter fiber tank at 29 °C and salinity 12 PPT, and observed for the embryonic stages until hatching took place. Organogenesis and developmental changes were recorded under a light microscope equipped with computer aided software (Nikon Eclipse E600). On 19^th^ day, most of the brooders released embryos which were pooled together in 20 liter fiber tank. Equal numbers of larvae were distributed among tanks with different experimental conditions (fig. 1). We chose following combinations of temperature and salinity: T*S [29°C/12 PPT] ,T_1_*S [31°C/12 PPT], T_2_*S [33.5°C/12 PPT], T*S1 [29°C/15 PPT], T_1_*S_1_ [31°C/15 PPT], T_2_*S_1_ [33.5°C/15 PPT], T*S_2_ [29°C/20 PPT], T_1_*S_2_ [31°C/20 PPT] and T_2_*S_2_[33.5°C/20 PPT]. The desired temperature in the tank was maintained using aquarium thermostat (Aqua Zonic, Singapore). The desired salinity was achieved by mixing fresh water with seawater of 35 PPT. The experimental tank consisted of 12 PPT sea water in glass aquaria with stock of 3540±82.76 larvae in it. During the experiment, larvae were fed with live *Artemia*.

**Figure 1:**
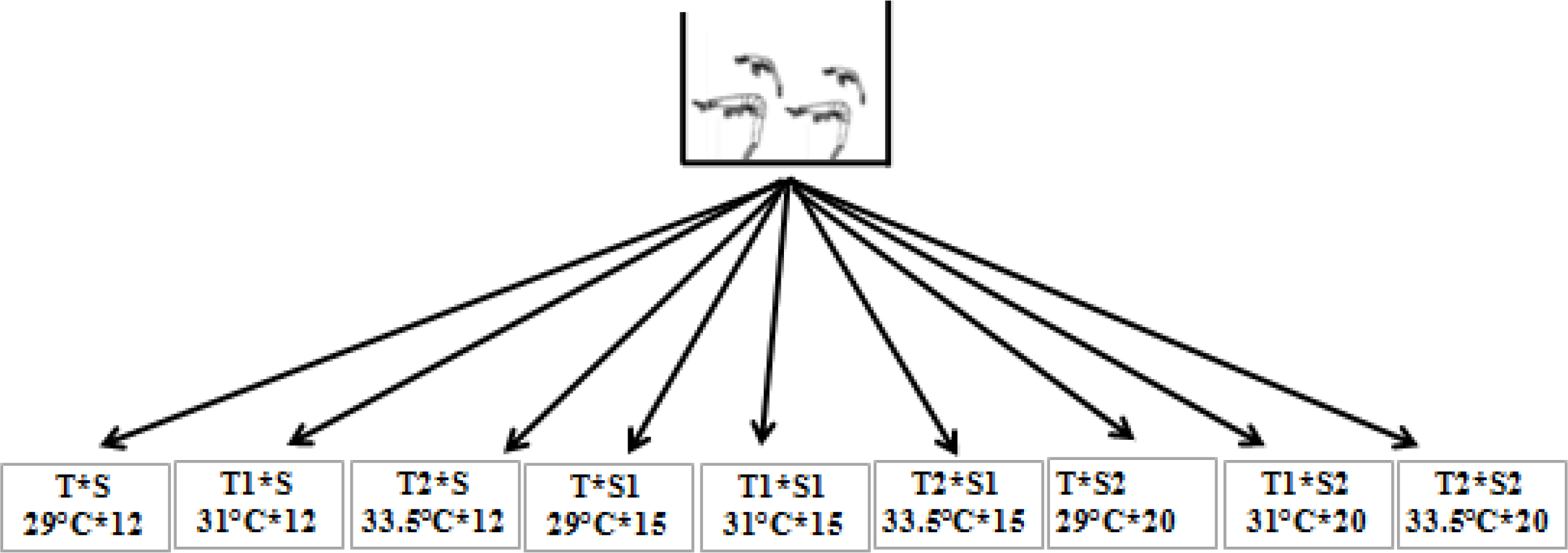
Experimental set up. The different experimental conditions. Where T*S [29°C/12 PPT] ,T_1_*S [31°C/12 PPT], T_2_*S [33.5°C/12 PPT], T*S_1_ [29°C/15 PPT], T_1_*S_1_ [31°C/15 PPT], T_2_*S_1_ [33.5°C/15 PPT], T*S_2_ [29°C/20 PPT], T_1_*S_2_ [31°C/20 PPT] and T_2_*S_2_[33.5°C/20 PPT]

The experiment lasted for 5 days from the day of hatching when most of the larvae were at 4^th^ stage. The yolk utilization were measured everyday till fully consumed. Following 5^th^ day, larval survival rate, growth rate, activity, and cardiac performance were measured.

### Yolk utilization

Depletion of yolk in larvae was determined by staining the live larvae with Nile Red. To quantify the yolk, 10 larvae from each treatment condition were stained with Nile Red (10 mg/ml) for 15 minutes, followed by washing using distilled water. The images were captured using the Epifluorescence microscope (Nikon Eclipse E600, excitation filter BP 490; barrier filter O515) equipped with digital camera and computer aided software (NIS-Elements). The images were analyzed by color threshold function in the image processing software Image J (Abràmoff et al., 2004). The total area and total intensity were calculated and represented in mm and pixels respectively. The representative image to show how we measured the area and intensity is shown in figure 2.

**Figure 2:**
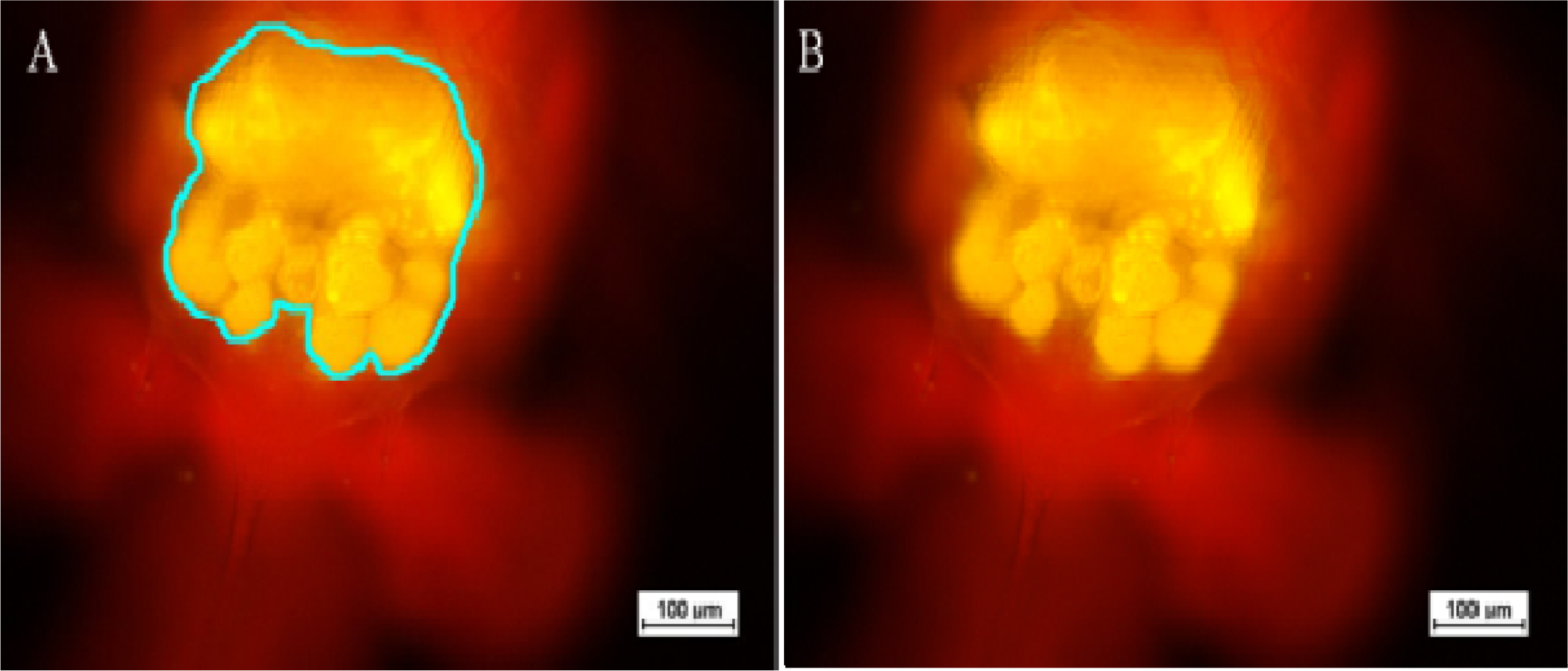
Yolk Utilization. Determination of depleted yolk volume in larvae treated under different conditions. A: Total yolk area marked by using image J; B- Total yolk Intensity analyzed by color threshold function by using image J

### Larval survival rate

Larval survival rate was estimated after 5 days of incubation at different conditions using the formulae:

Larval survival rate (%) = {initial number - (initial number - final number)/initial number}*100.

### Morphometrics of larvae

Morphometric analyses were conducted on the images taken by stereomicroscope equipped (Motic (Xiamen) Electric Group Co., Ltd, China) with digital camera and computer aided software (Motic image plus 3.0). Total length was estimated after 5 days of incubation by subtracting initial length from final length and represented in millimeter (mm). The representative image for the total length is shown in figure 3.

**Figure 3:**
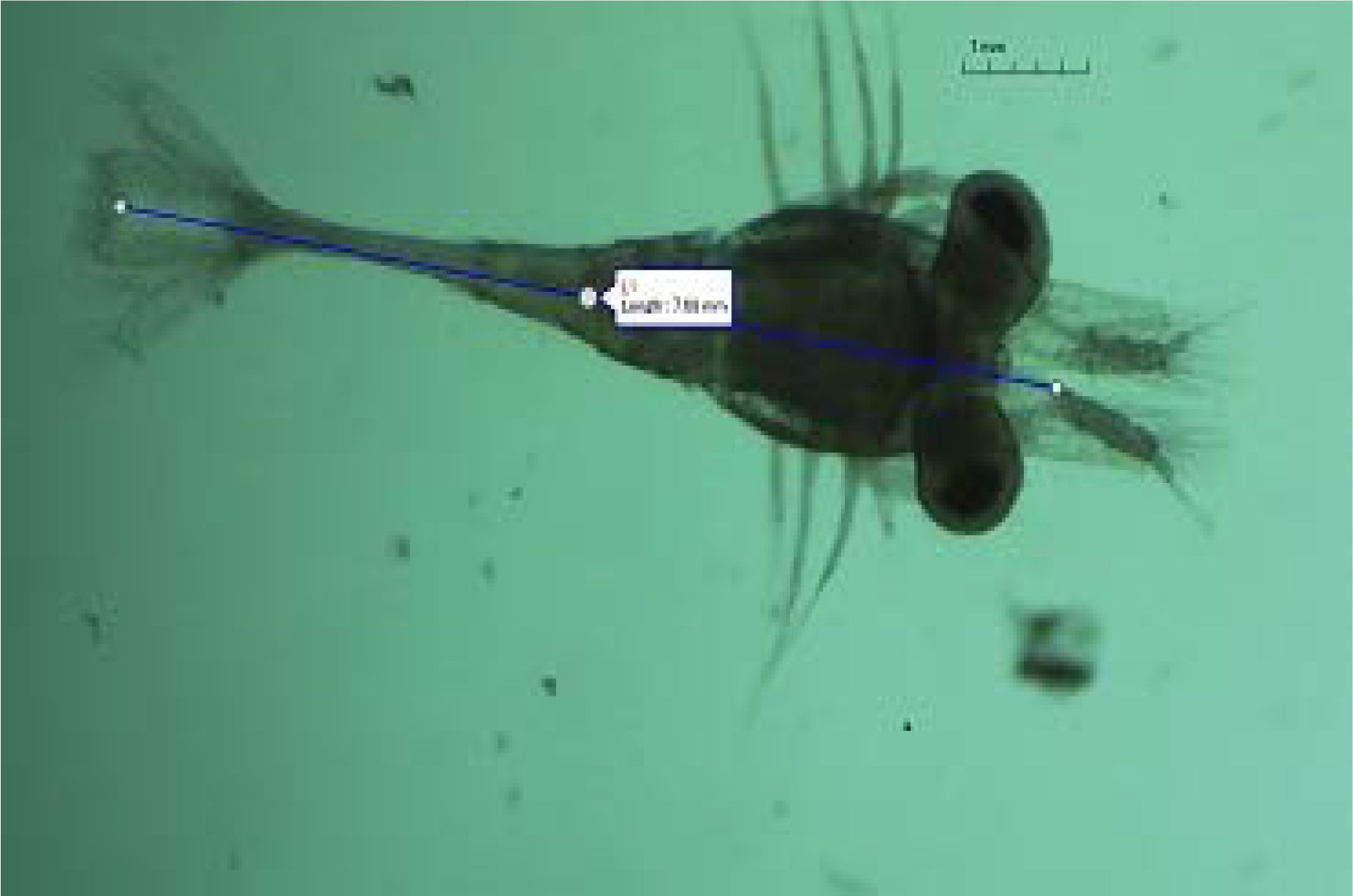
Morphometrics. Total length measured by using stereo dissection microscope

### Larval activity

Larval activity, defined as the rate of maxilliped movement, was determined in the video taken using stereomicroscope (Ceballos-Osuna et al., 2013). Larva was trapped in a drop of water on cavity slide and covered with a cover slip. The water drop was taken from the respective experimental tank, and video was recorded at 5X magnification using stereomicroscope (Motic (Xiamen) Electric Group Co., Ltd, China) for 2 minutes. Videos were parsed into 10s segments (free video cutter v 10.4), and slowed to 25% from original speed (VLC media player V. 2.4.4) for counting maxilliped movements (first three feeding legs) for at least 3 individuals. The results were represented in beats per minute (bpm).

### Cardiac performance

Heart rate (ƒ_H_) and stroke volumes (V_S_) were determined from the same video recorded for the larval activity. The videos were slowed down to 25% from original speed (VLC media player V. 2.4.4) and zoomed to count accurate ƒ_H_. A representative video showing the heart beat and maxilliped movements can be seen in supplementary file video 1. Screen marker (Epic pen V. 3.0) was used to mark end-diastolic and end-systolic perimeter. V_S_ were determined by calculating the difference between the end-diastolic volume (EDV) and end-systolic volume (ESV), assessed using Image J.

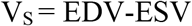

EDV and ESV were assumed as prolate spheroids (Harper and Reiber, 2004; Storch et al., 2009), so following equation was used to calculate volume.

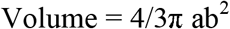

Where, a is the radius of major diameter and b is the radius of minor diameter (from image analysis). Individual cardiac output (Q) was determined as a product of V_S_ and *f*_H_. The representative images showing EDV and ESV and VS are given in figure-4.

**Figure 4:**
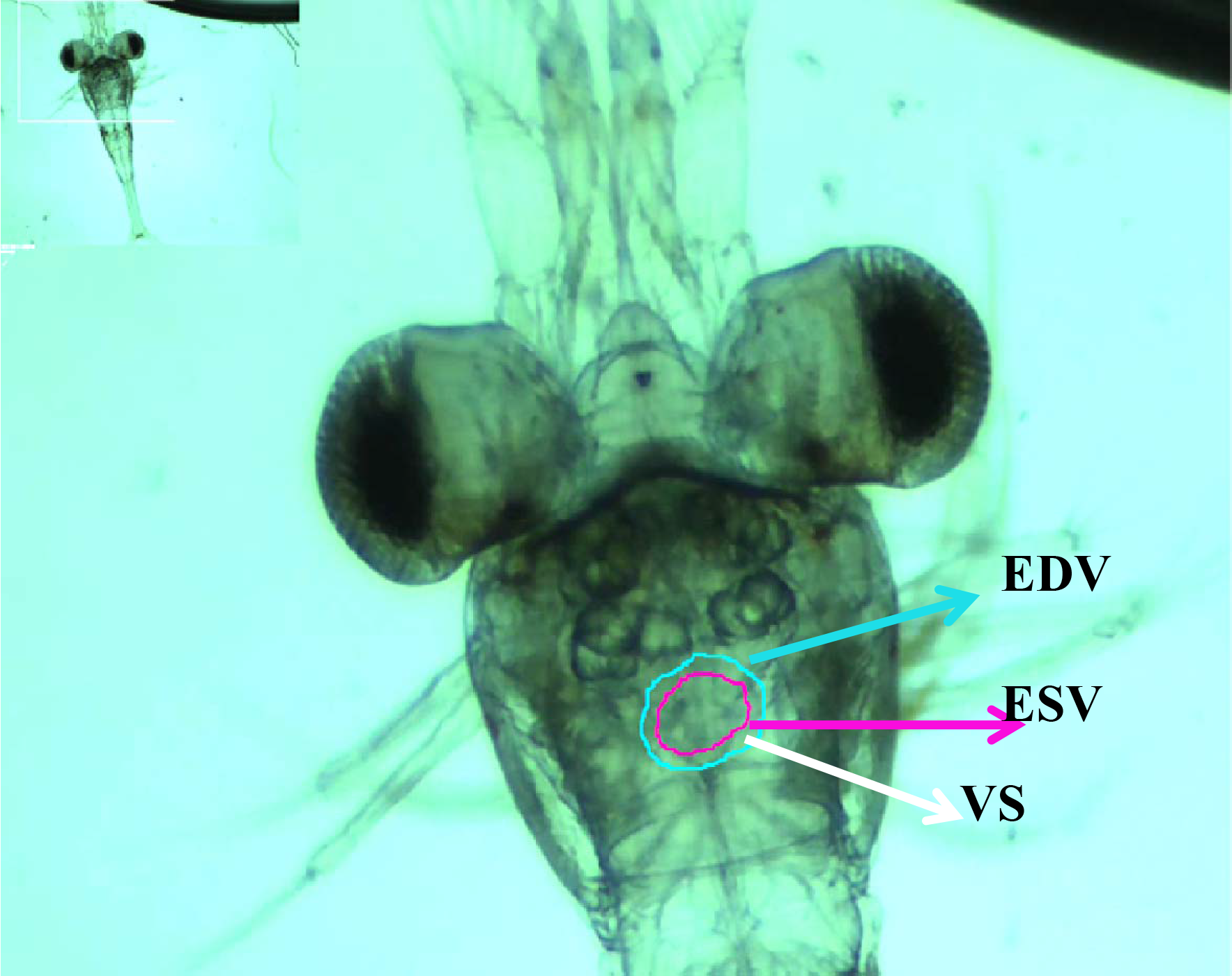
Cardiac performance. A marked picture of larval heart showed EDV (maximal area) and ESV (minimal area). V_s_ was calculated as the difference between EDV and ESV

## Statistical analysis

Data were tested for normality and homogeneity using Shapiro–Wilk and variance test. After successful completion of these parameters, we performed two way ANOVA and post hoc tukey’s tests on the mean values for finding the independent and dependant effect of temperature and salinity. All the statistical analyses were performed using SPSS v. 22 (Corp, 2013).

## Results

### Larval survival rate

Immediately after hatching, we collected the larvae and concentrated in 12 PPT seawater with a density of 71±3.7 individuals ml^-1^. Soon after, equal volume of water assuming equal numbers of larvae (approx. 3500 ± 82.76 individuals) were distributed among all the experimental conditions. On day 5^th^, we observed 100% mortality at the higher temperature conditions (33.5 ± 0.5 °C) in all the salinity conditions (12 PPT, 15 PPT, and 20 PPT). However, among other experimental conditions, the survival rate varied between 76-96 % (Table 1). Unfortunately, due to technical faults, we lost all the larvae in tank with 20 PPT and 31°C on day 1^st^. At the ambient temperature of 29 °C and different salinity, larvae survival rates were 96 %, while at 31°C; 12 PPT and 31°C; 15 PPT showed 86% and 76% survival rate respectively.

**Table 1.**
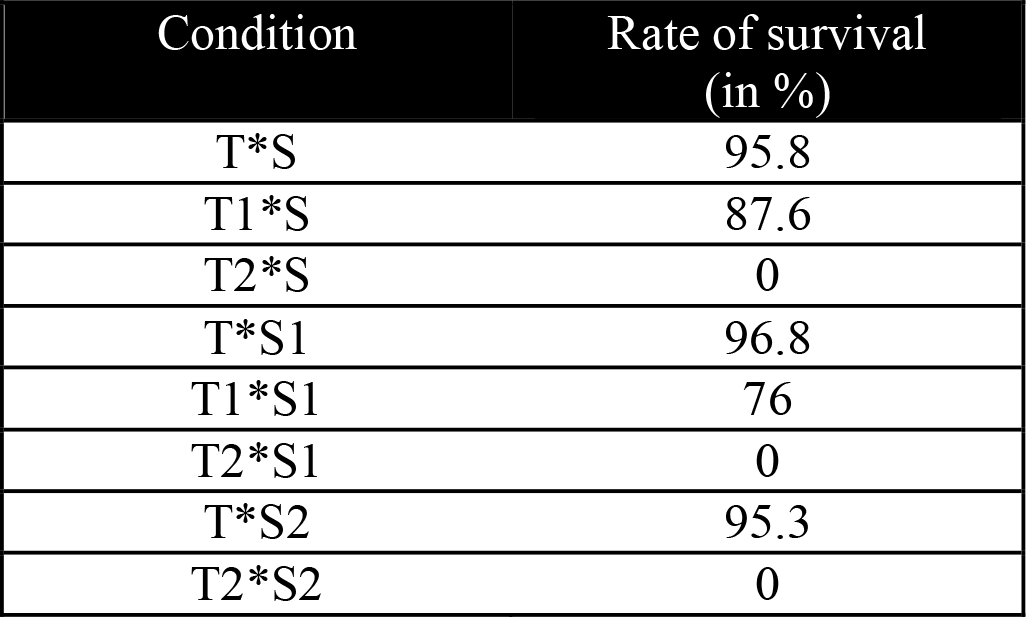
Survival rate of larvae exposed under different conditions

### Yolk consumption

Post-embryonic yolk was depleted at different rates under varying experimental conditions. In the ambient condition (12 PPT; 29 °C), yolk lasted till day 4^th^. Moderate increase in temperature (31 °C) and salinity alone did not show any effect on the rate of yolk consumption, whereas higher temperature (33.5 °C), alone and in combination with increased salinity caused faster depletion of yolk. In these cases, the yolk was almost depleted either on day 2^nd^ or day 3^rd^ (fig. 5). The utilization of the yolk under different conditions in terms of total area and intensity has been shown in figure 6.

**Figure 5:**
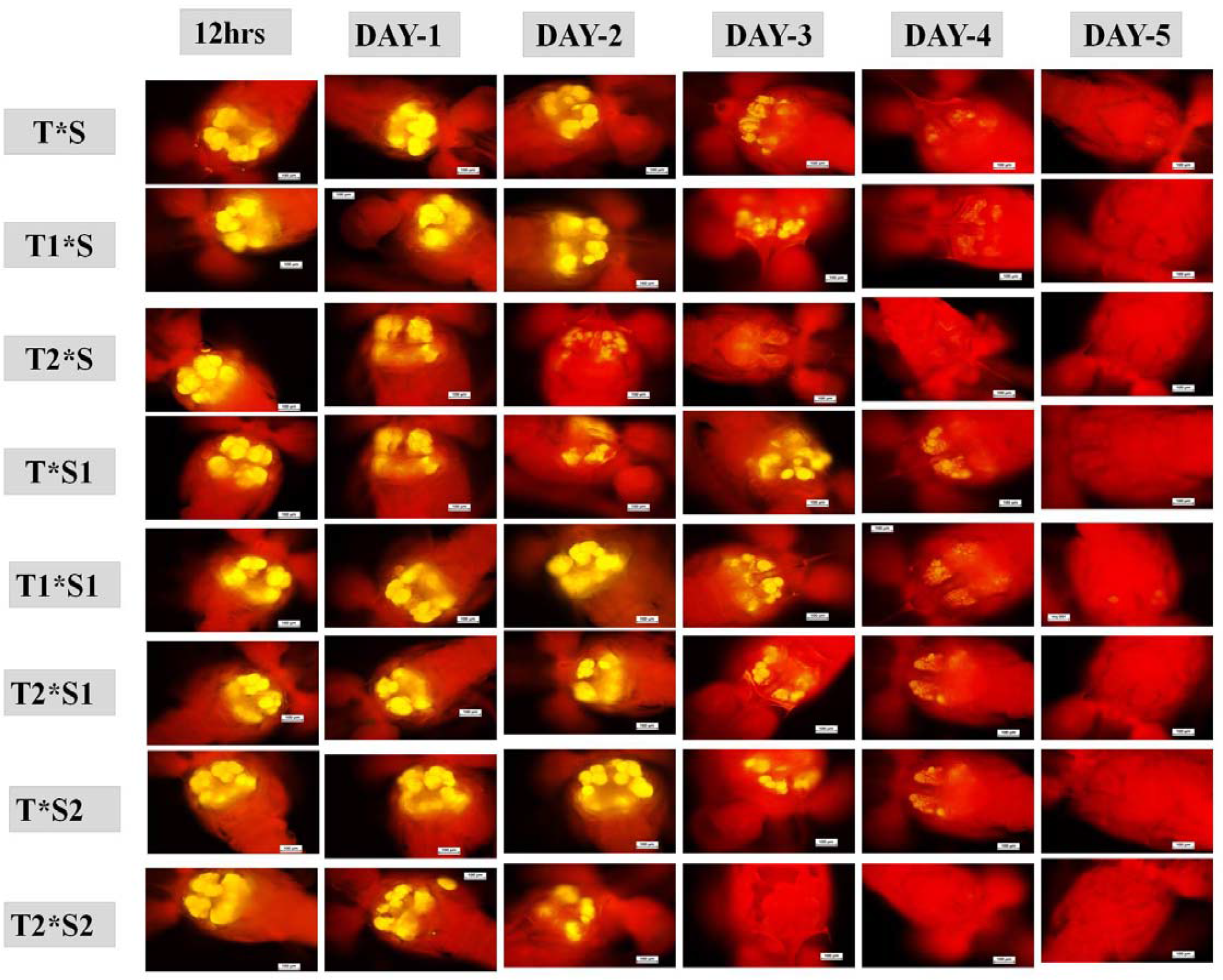
Utilization of yolk. Depletion of yolk in different conditions T*S-29°C/12 PPT; T_1_*S-31°C/12 PPT; T_2_*S-33.5°C/12 PPT; T*S_1_-29°C/15 PPT; T_1_*S_1_-31°C/15 PPT; T_2_*S_1_-33.5°C/15 PPT; T*S_2_-29°C/20 PPT; T_2_*S_2_-33.5°C/20 PPT

**Figure 6:**
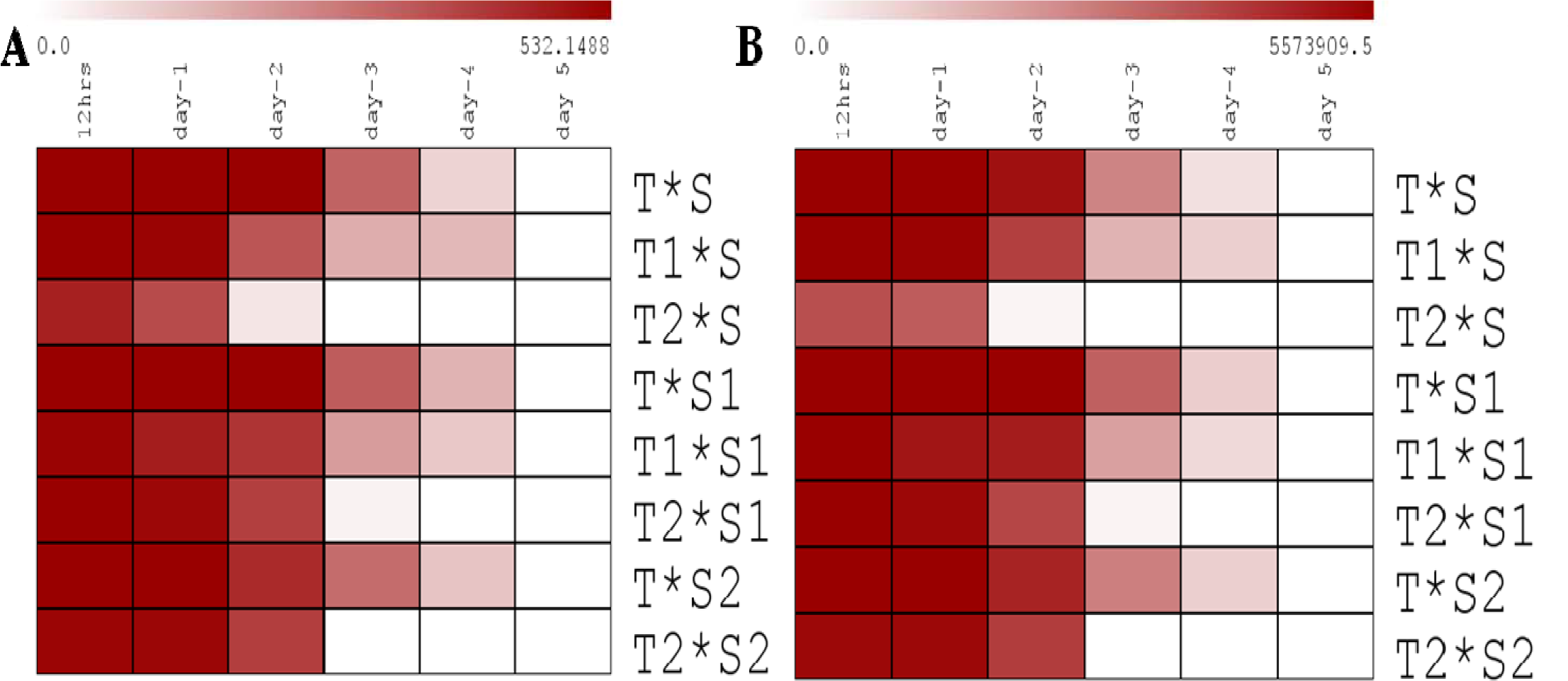
Yolk utilization in total area and intensity. **A**- Total yolk area in mm and B- Total yolk Intensity in pixel T*S-29°C/12 PPT;T_1_*S-31°C/12 PPT;T_2_*S-33.5°C/12 PPT; T*S_1_-29°C/15 PPT;T_1_*S_1_-31°C/15 PPT;T_2_*S_1_-33.5°C/15 PPT;T*S_2_-29°C/20 PPT;T_2_*S_2_-33.5°C/20 PPT

### Morphometrics of larvae

Larval morphometrics were analyzed by measuring total length after 4^th^ day (fig. 7). In general, we observed an increase in the total length of larvae upon exposure to higher temperature and salinity independently or in combination up to 4^th^ day (*p<0.001(temperature); p<0.01(salinity); p<0.05(temperature*salinity)*), however the larvae in the higher temperature (33.5 °C) in all three salinity (12, 15, and 20 PPT) conditions died on 5^th^ day.

**Figure 7:**
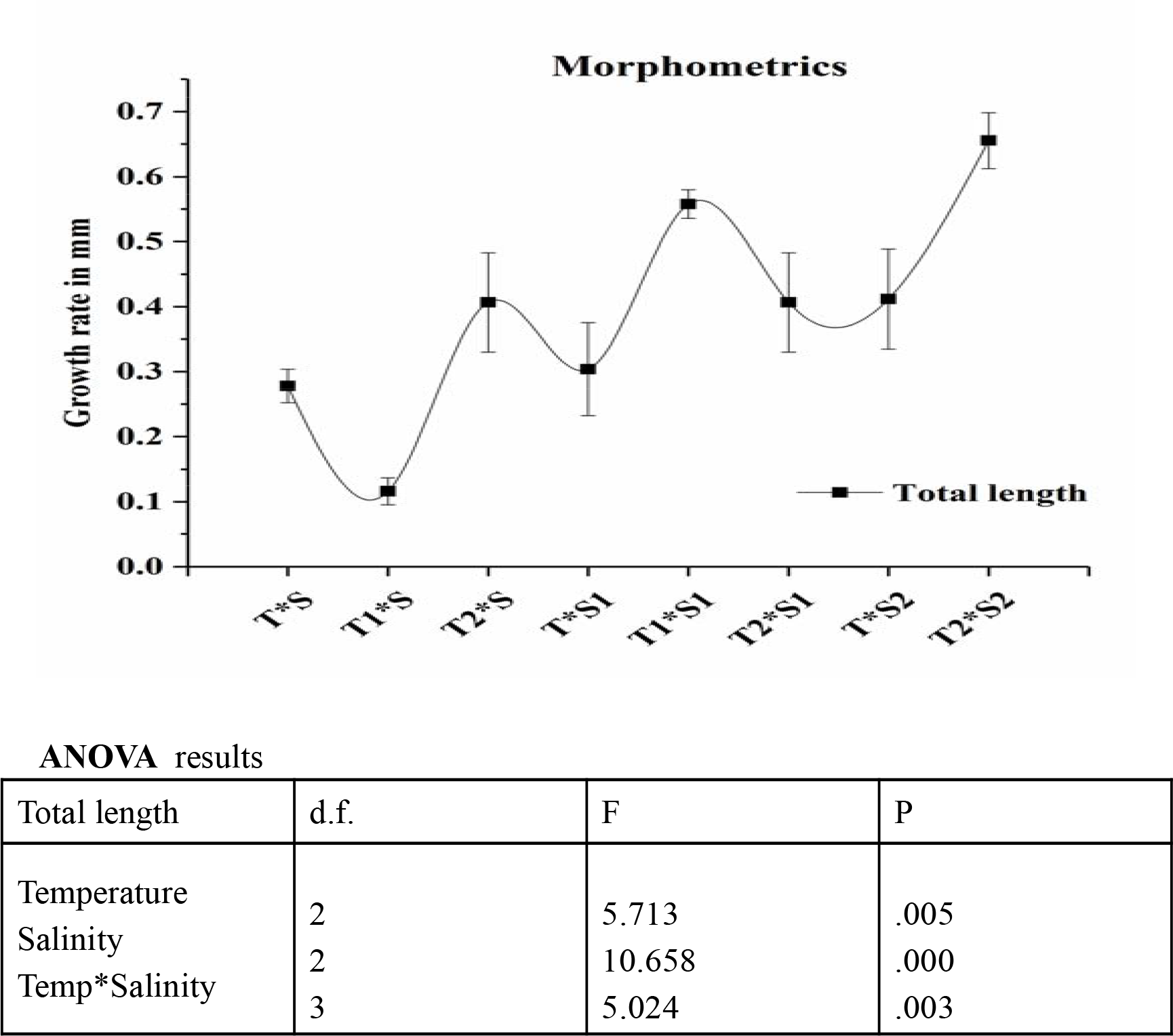
Growth rate of larvae. Growth rate of larvae on 4th day was calculated by subtracting the growth in mm on 1st day from 3rd day. Where T*S-29°C/12 PPT] ,T_1_*S-31°C/12 PPT; T_2_*S-33.5°C/12 PPT; T*S_1_-29°C/15 PPT; T_1_*S_1_-31°C/15 PPT; T_2_*S_1_-33.5°C/15 PPT; T*S_2_-29°C/20 PPT; T_2_*S_2_-33.5°C/20 PPT

### Larval activity

Larval activity was measured in terms of maxilliped movement which ranged from 90 to 250 bpm across different conditions (fig. 8). Mean maxilliped frequency decreased significantly at higher temperature and salinity conditions than the ambient conditions (*p<0.05(temperature); p<0.001(salinity)*, however, the interaction of temperature and salinity did not show any effect on them (p = 0.760).

**Figure 8:**
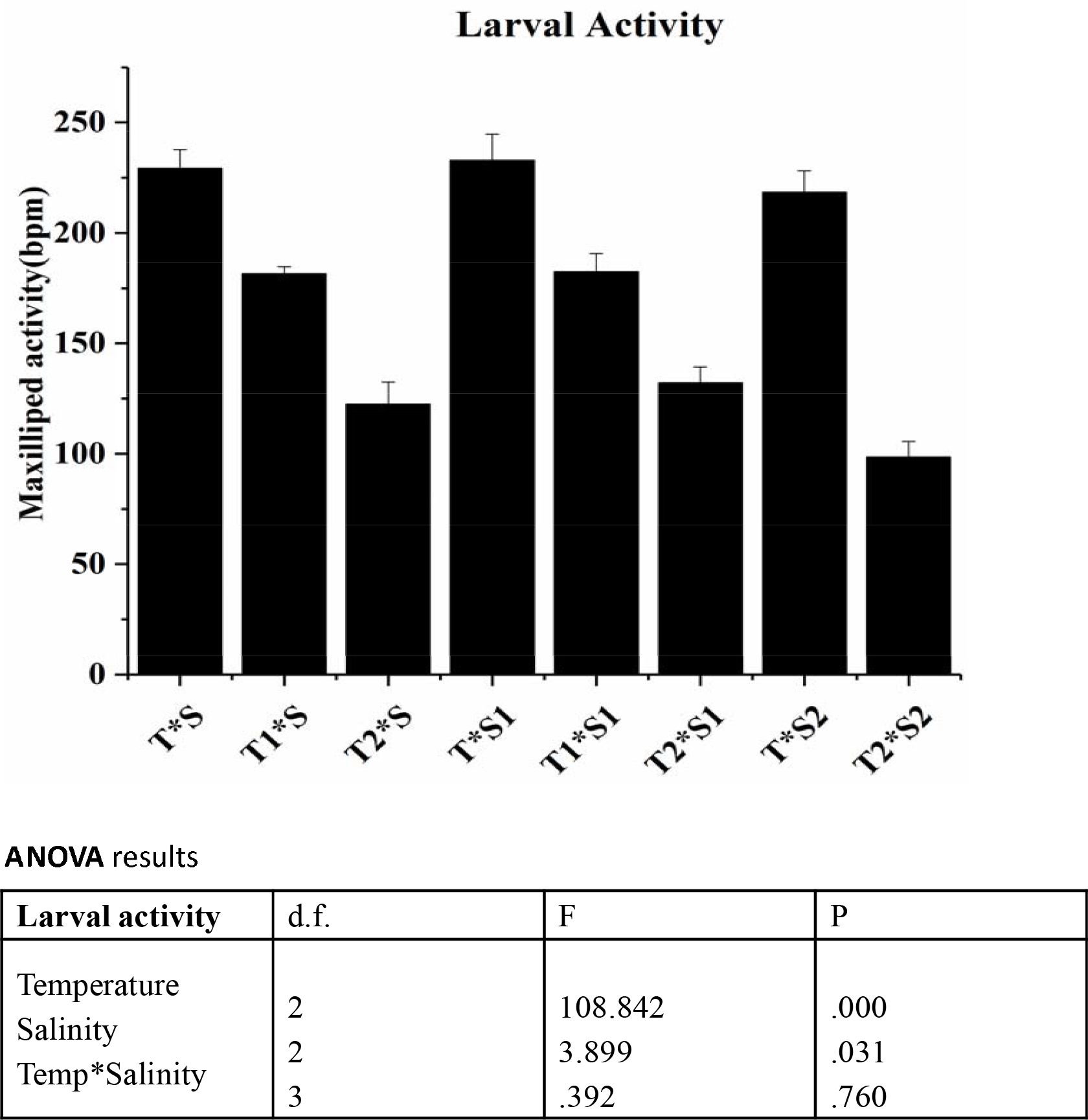
Larval activity. Maxilliped activity of larvae on 4th day under different conditions. Where, T*S-29°C/12 PPT; T_1_*S-31°C/12 PPT; T_2_*S-33.5°C/12 PPT; T*S_1_-29°C/15 PPT; T_1_*S_1_-31°C/15 PPT; T_2_*S_1_-33.5°C/15 PPT;T*S_2_-29°C/20 PPT;T_2_*S_2_-33.5°C/20 PPT

### Cardiac performance of larvae

Mean cardiac performance in larval stage was significantly affected under the treated conditions. We found larval heart beat (*f*_H_) in the range of 200-300 beats min^-1^, with maximum beating in higher temperature and salinity conditions (20 ppt; 33.5 °C) and minimum in ambient condition 12 ppt; 29°C. Temperature, salinity, and their interactions showed significant effect on *f*_H_ (*p<0.001(temperature); p<0.01(salinity); p<0.05(temperature*salinity))*. Larval stroke volumes (V_s_) across different conditions were in the range of 0.005-0.020 nl beats^-1^, higher in ambient conditions and lowest in higher temperature and salinity conditions (fig. 9). Temperature, salinity, and their interactions showed significant effect on *V*_s_ (*p<0.001(temperature); p< 0.001 (salinity); p<0.05(temperature*salinity))*. Cardiac output (Q□) which is the product of *V*_s_ * *f*_H_ was in the range of 2-5 nl min^-1^, with higher in the ambient condition and lowest in the higher temperature and salinity conditions. Temperature and salinity independently showed significant effect on Q□ *(p<0.05(temperature); p<0.001 (salinity))*, while their interaction seemed not to have any impact on Q□ *(p=0.099)*. *V*_s_ and Q□ were following similar patterns in wave manner; both of them were inversely related to fH across different treatment conditions (figure 9).

**Figure 9:**
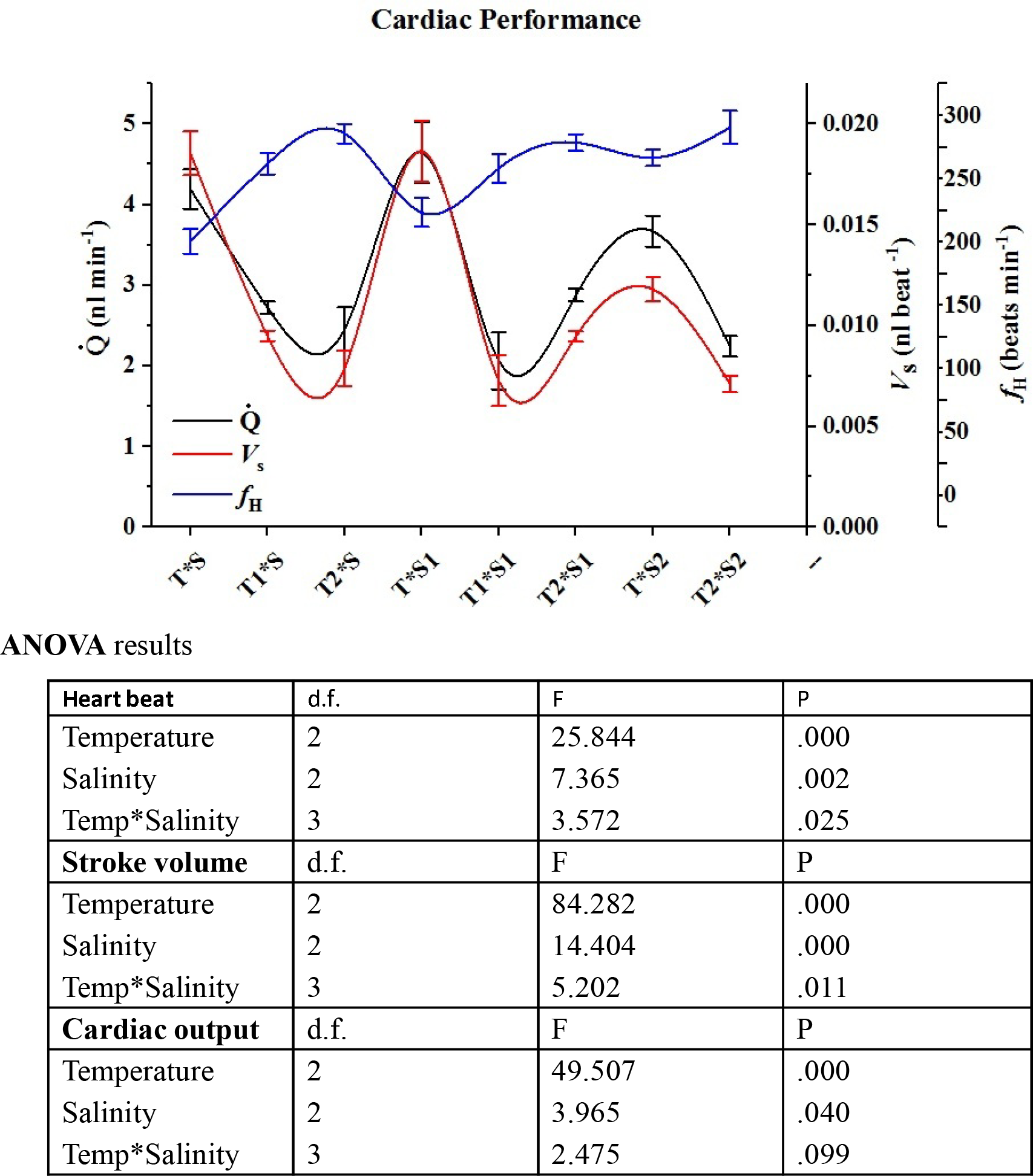
Cardiac performance. Cardiac performance of larvae on 4^th^ day. Where T*S-29°C/12 PPT; T_1_*S-31°C/12 PPT; T_2_*S-33.5°C/12 PPT; T*S_1_-29°C/15 PPT; T_1_*S_1_-31°C/15 PPT; T_2_*S_1_-33.5°C/15 PPT; T*S_2_-29°C/20 PPT; T_2_*S_2_-33.5°C/20 PPT

## Discussion

In this study, we assessed physiological performance of *M. rosenbergii* early life stages following exposure to varying temperature and salinity to understand their response to climate change stress conditions. Physiological performances are discussed in two broad categories: 1) survival and growth; 2) larval activity and metabolic performance. Overall, we find a small rise in temperature and salinity may result in sub lethal physiological rate reductions in *M. rosenbergii* early life stages, but the substantial increase in temperature and salinity may be detrimental.

## Survival and growth

Salinity and temperature are the important environmental factors affecting survival, growth and distribution of many aquatic organisms (Habashy and Hassan 2010; Kumlu et al. 2000). In the adult *M. rosenbergii* the survival rate varied between 91% (at 0 PPT) and 78% (at 20 PPT), and the prawn exhibited lowest final average weight at 20 ppt seawater and higher at 10 PPT salinity (Chand et al., 2015). Similarly, Habashy et al (2010) reared the juvenile prawns for eight months in different salinity and temperature conditions revealing that growth of the prawn increased as temperature increased from 24 to 29 °C, but declined at the higher temperature (34 °C). Also, with increase in salinity from 0 to 16 PPT, growth of female prawn decreased at all temperatures tested (Habashy and Hassan, 2010). Recently, Mohanty et al. (2016), conducted survival experiments on zoeae and post larvae of *M. rosenbergii* for combined effects of salinity and temperature, in which, post larvae showed maximum survival at 31 °C which declined both at lower and higher temperature of 25 °C and 35 °C, respectively. In the line of these previous results on adults and post larvae, we also found that larval survival rates were higher at optimum temperature of 29 °C in 12 PPT, 15 PPT, and 20 PPT. However, survival rate decreased with increase in temperature at all tested salinity conditions. For the higher temperature conditions (33.5 ± 0.5 °C) with all salinity conditions (12 PPT, 15 PPT and 20 PPT) no survival was recorded. Similar to our results, Mohanty et al. (2016) reported lower survival rate for *M.rosenbergii* zoeae (Z1-Z5) at 35 °C and 15-18 PPT salinity.

For larval growth, there was an increase in total length of larvae up to 4^th^ day for salinity (p<0.01), temperature and combined temperature and salinity (p<0.05). Although temperature was major determining factor for growth (Kumlu et al., 2000), the quality of larvae was compromised and died in 5^th^ day for the higher temperature (33.5°C) in all salinity conditions.

### Larval activity and metabolic performance

Lipid in yolk acts as energy sources for the early stages of larvae, ensuring the first successful molt and supporting the survival of early larvae before it started feeding (Yao et al., 2006). The effects of temperature on metabolic and developmental rates are expressed through changes in the consumption speed of reserves (García-Guerrero, 2010). In the present study, larval yolk in both ambient temperature and 31 °C lasted for 4 days, but for the higher temperature, the yolk depletion was completed either on day 2^nd^ or day 3^rd^ (figure 5, 6). Under the increased temperature, the rate of metabolic processes hiked, demanding more energy, and causes the early depletion of energy reserve. The influence of temperature on the utilization of yolk content was reported in several aquatic species ((Evjemo et al., 2001; García-Guerrero, 2010; Holland, 1978).

During the stressful condition, organisms try to maintain its homeostasis. The capacity of controlling cardiovascular function is one such function to maintain the organism’s oxygen consumption rate and activity of the organism (Ern et al., 2015). Temperature alters cardiac performance in crustaceans, as has been reported in several other organisms (Goudkamp et al., 2004; Jury and Watson, 2000; Morris and Taylor, 1985). Salinity also affects the sensitivity of organism; thereby affecting their oxygen consumption (Barton and Barton, 1987). In the present study, both salinity and temperature are shown to influence stroke volume and cardiac output, ultimately affecting the oxygen transport capacity of the animal. Larval heart beat (*f*_H_) in *M. rosenbergii* larvae increased significantly with elevated temperature and salinity, but at the same time, stroke volume (V_s_) decreased, that reduced cardiac output (Q□) as well as oxygen transport capacity. Ern et al. (2014) showed that heart rates and ventilation rates increased and stroke volume decreased with increasing temperature in adult *M. rosenbergii*. They also showed that the animals retained their 76% of aerobic scope at 30°C. We observed the lower stroke volume and cardiac output in the present study, which may have affected the oxygen consumption rate of organism. Hence, the temperature and salinity beyond tolerance level could initiate anaerobic metabolism with detrimental effects.

Further, the decrease in aerobic scope could affect the function and behavior of larval activity in order to maintain pejus temperatures for long term survival (Portner and Knust, 2007; Wang and Overgaard, 2007). For example, the rise of temperature above 15°C in the kelp crab *Taliepus dentatus* constrained the aerobic scope and affected the level of maxilliped activity (Storch et al., 2009). This was observed in the present study as well, in which the mean maxilliped frequency was significantly lowered in elevated temperature and salinity conditions. However, under the combined conditions of salinity and temperature, larval activity did not show significant effect as similar to our cardiac output results.

Corollary, this study shows that substantial increase of temperature and salinity may result in negative impact on the survival, growth, cardiac performance and activity of the early life stages of *M. rosenbergii*. The effect of climate change stressors thus could restrain the tolerance capability and physical fitness of the early life stages of this freshwater prawn, thereby affecting the successful persistence of the population.

## Acknowledgments

The authors are thankful to Dr. T. Sasipraba, the Pro-Vice Chancellor, for her constant support in establishing aquaculture facility without which work would not have been possible. We deeply thank Dr. V. Balasubramanian for his assistance in sample collection and Dhanalekshmi hatchery for providing prawns. We thank Dr. Vinitha vishwakarma for letting us use Epifluorescence microscope.

## Author contributions

VSS, JJ conceptualized the work with assistance from AK, VE, TS and SP. VSS, JJ, AK, PR, and VE performed sampling campaign and laboratory experiments. JJ analyzed the results and VSS, AK, TS, VE, UK helped her in interpreting results. VSS and AK wrote the first draft which was subsequently corrected by all authors.

